# Model-based evaluation of Targeted-Antibacterial-Plasmids (TAPs) transfer kinetics and resensitization of pOXA-48 carbapenem-resistant *Escherichia coli*

**DOI:** 10.64898/2026.09.01.748484

**Authors:** Sophie Marolleau, Jérémy Moreau, Julien M. Buyck, Sarah Bigot, Christian Lesterlin, Nicolas Grégoire, Vincent Aranzana-Climent

## Abstract

**Background:** Targeted-Antibacterial Plasmids (TAPs) are engineered mobile genetic elements that use bacterial conjugation to deliver selective CRISPR/Cas9 antibacterial activity against a specific target strain. Yet, the efficiency of TAPs is typically evaluated at a single time point, whereas the success of TAP-mediated resensitization critically depends on the dynamics of plasmid transfer and the complex interactions between bacterial subpopulations. This is the first study to evaluate the efficiency of a conjugation-based antibacterial approach at the subpopulation level, using an analytical framework analogous to that used for conventional antibiotics. Here, we investigate which process limits resensitization by TAP_F_-dCas9-OXA48: plasmid delivery, dCas9 activity, or the emergence of refractory and escape populations.

**Methods:** We fitted a mechanistic model of five interacting subpopulations (donors, recipients, transconjugants, escapers, and recusants) to 44 longitudinal conjugation experiments and used the fitted model to explore a range of biologically relevant scenarios.

**Results:** Using longitudinal conjugation data spanning 24 h, we show that up to 24% of recipients become recusants within 24h, refractory to further conjugation *via* entry exclusion, while secondary transconjugant emergence stays below 0.01%. Overall resensitization efficiency reaches up to 80%.

**Conclusion:** Plasmid transfer, rather than dCas9 repression, therefore appears to be the main bottleneck limiting the efficiency of TAP_F_-dCas9-OXA48 efficiency. These results identify plasmid delivery as a key engineering target for improving the performance of future TAPs.

## Introduction

Antibiotic resistance represents a major global health challenge limiting the effectiveness of current treatments and increasing the need for alternatives to conventional antibiotics (1). Among the most critical threats, carbapenem-resistant *Escherichia coli* has been classified as a critical priority pathogen by the World Health Organization, reflecting its high mortality burden, increasing resistance trends, and severely limited therapeutic options (2,3). Carbapenem resistance in *E. coli* is largely mediated by the acquisition of carbapenemase-encoding genes, such as *bla*_OXA-48_, carried by the conjugative plasmid pOXA-48 (4). While horizontal gene transfer (HGT) *via* conjugation drives the dissemination of such resistance determinants, it can also be repurposed as a delivery platform/system for engineered genetic tools into bacterial populations.

In this context, targeted antibacterial plasmids (TAPs) have recently emerged as a promising strategy to selectively eliminate or reprogram antibiotic-resistant bacteria (5–7). Reuter *et al*. (5) demonstrated that mobilisable plasmids encoding CRISPR/Cas9 systems could achieve strain-specific antibacterial activity by targeting resistance genes, leading to selective killing or plasmid curing. Building on this concept, Djermoun *et al*. (2023) (6) expanded the applicability of TAPs by reprogramming them to function across a broader range of bacterial hosts, thereby widening the restricted host range of earlier designs. More recently, Derollez *et al*. (2026) (7) demonstrated the potential clinical relevance of this approach by achieving efficient and specific targeting of pathogenic *E. coli* strains harboring the *bla*_CTX-M-15_ β-lactamase gene, resulting in targeted killing and resensitization to β-lactam antibiotics. Together, these studies establish TAPs as versatile and programmable antimicrobial tools capable of precise targeting and modulation of resistant bacterial populations.

TAPs can be engineered to carry either CRISPR/Cas9 or CRISPR/dCas9 systems. CRISPR/Cas9 approaches typically rely on the introduction of sequence-specific double-strand breaks/DNA cleavage, leading to target-cell killing or elimination of resistance plasmid and, consequently, restoration of antibiotic susceptibility. In contrast, CRISPR/dCas9 systems employ a catalytically inactive Cas9 that binds to target genes without cleavage, thereby repressing their expression through transcriptional silencing (8). Both strategies enable selective targeting of resistant subpopulations while limiting effects on non-target bacteria, thereby offering a controlled and precise ecological intervention. However, the efficiency of TAP-mediated resensitization critically depends on the dynamics of plasmid transfer and the complex interactions between bacterial subpopulations.

Consequently, TAP-mediated resensitization emerges from a sequence of biological processes, including plasmid transfer, CRISPR targeting, and the potential emergence of subpopulations that escape TAP system or become refractory to further conjugation. Distinguishing the contribution of each of these processes is essential for identifying the factors that ultimately limit antibacterial efficiency. Understanding and quantitatively describing these conjugation-based antimicrobial strategies therefore requires integrating bacterial population dynamics, plasmid transfer, and antimicrobial activity within a unified framework. Such an approach is conceptually analogous to pharmacokinetic/pharmacodynamic (PK/PD) and time-kill curve (TKC) analyses used to quantify the effects of conventional antibiotics. Applying similar quantitative principles to engineered conjugative systems could therefore enable TAPs to be characterized as dynamic antimicrobial interventions rather than solely as genetic delivery tools.

Mathematical models have previously been developed to describe plasmid transfer dynamics, often relying on mass-action formulations to capture conjugation processes (9,10). These models provide a useful framework to quantify transfer rates and predict population dynamics over time. However, existing models have not yet been applied to engineered conjugative systems delivering CRISPR-based antimicrobials. In particular, the relative contributions of plasmid transfer, CRISPR activity, and the emergence of refractory or escape subpopulations to overall resensitization remain unclear. In this study, we investigated the dynamics of delivery *via* conjugation of a CRISPR/dCas9-based TAP system targeting *bla*_OXA-48_ (TAP_F_-dCas9-OXA48), using longitudinal conjugation experiments coupled with a mechanistic model of interacting with bacterial subpopulations.

## Results

### 1. Experiments

#### 1.1 Counting bacterial subpopulations

The *bla*_OXA-48_ gene carried on pOXA-48 confers resistance to carbapenems, the clinically relevant phenotype associated with this plasmid. In this study, ampicillin resistance was used as a phenotypic readout of *bla*_OXA-48_ expression, allowing selection and discrimination of bacterial subpopulations on agar plates.

TAPs are mobilizable plasmids whose conjugative transfer depends on a helper plasmid that provides the machinery required for mating-pair formation and DNA transfer and encodes entry-exclusion functions. In the present system, the helper plasmid confers tetracycline resistance, enabling helper-carrying bacteria to be identified on tetracycline-containing agar plates.

Experimental time-course data were collected and used to develop a mathematical model describing the evolution of the different bacterial subpopulations involved in the process, namely donors, recipients, transconjugants, escapers, and recusants (Figure 1). Donors carry both the TAP and the helper plasmid, while recipients carry pOXA-48 alone. Conjugation between these two populations can generate three distinct recipient-derived subpopulations depending on which plasmids are acquired and whether TAP-mediated repression is effective. Transconjugants acquire all three plasmids; in these cells, the TAP successfully represses blaOXA-48 expression, resulting in restoration of ampicillin susceptibility. Escapers carry the same three plasmids, but remain ampicillin-resistant, indicating failure of TAP-mediated blaOXA-48 repression. Finally, recusants carry pOXA-48 and the helper plasmid, but not the TAP. Three possible origins were considered for recusants: acquisition of the helper plasmid alone from donors, acquisition of the helper plasmid alone from transconjugants acting as secondary donors, or loss of the TAP following transconjugant formation. Only the first and third routes — helper acquisition from donors, and TAP loss by transconjugants — were supported by the data and retained in the model, while helper transfer from transconjugants was excluded.

**Figure 1.**
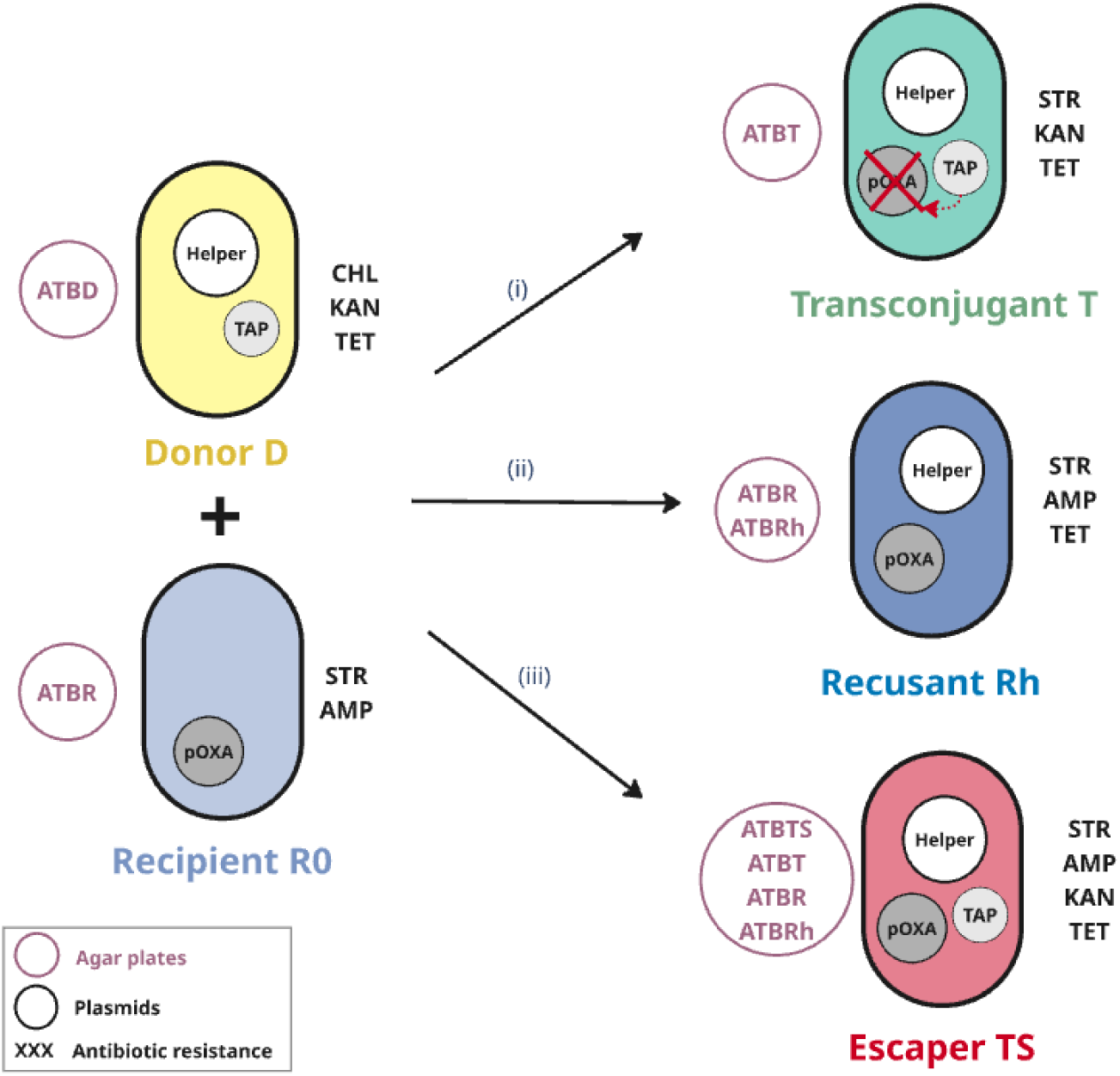
Outcomes of conjugative transfer between the TAP donor and the pOXA-48-carrying recipient. The donor strain (D) carries two plasmids: the helper plasmid, which encodes the conjugation machinery, and the TAP (Targeted-Antibacterial Plasmid), which is mobilized in trans via the helper. The recipient strain (R0) carries pOXA-48, the plasmid responsible for ampicillin resistance. Following mating, three main outcomes can be observed, depending on which elements are transferred and whether the TAP is functionally effective in the new host: **(i) Transconjugants (T):** Both the helper and the TAP are transferred to the recipient, and the TAP successfully represses expression of the carbapenemase gene (bla_OXA-48_) carried by pOXA-48. The transconjugant is resensitized to ampicillin. **(ii) Recusant (Rh):** Only the helper plasmid is transferred, without the TAP. The recusant is refractory to further conjugation (entry exclusion) and remains ampicillin resistant. **(iii) Escaper (TS):** The recipient receives both the helper and the TAP, but bla_OXA-48_ expression is not repressed. The escaper remains ampicillin-resistant despite acquiring the TAP. Purple circles indicate the selective agars used to isolate each strain type. Antibiotic resistance phenotypes of each strain are shown in black (CHL: chloramphenicol; TET: tetracycline; STR: streptomycin; AMP: ampicillin; KAN: kanamycin).

Subpopulation identification after conjugation experiments was achieved by phenotypic selection using antibiotic-containing agar plates, and multiplex PCR confirmed that this phenotypic discrimination was fully concordant with genotypic characterization (Supp. material S1). Donors (D) were found to carry both the TAP_F_-dCas9-OXA48 and the F-Tn10 helper plasmid; recipients (R0) harbored pOXA-48. Among subpopulations that appear during conjugation, both Transconjugants (T) and escapers (TS) carried pOXA-48, the TAP and the helper plasmids; escapers were distinguished from transconjugants by their resistance to ampicillin despite the presence of the TAP. Recusants (Rh) harbored pOXA-48 and the helper plasmid only, consistent with their refractory status (entry exclusion) to further conjugation. Taken together, these results confirm that phenotypic and genotypic characterizations are fully concordant across all defined subpopulations, thereby validating the analytical approach and experimental procedure used to identify and monitor these subpopulations throughout the study.

Raw experimental data consisted of bacterial counts obtained from antibiotic-containing agar plates (ATBD, ATBR, ATBRh, ATBT, and ATBS). Table 2 lists the selective agar plates used, along with the subpopulations they select for. Since some plates select for more than one subpopulation at once, these raw counts were used directly as model inputs, with subpopulation abundances obtained as model outputs through the appropriate subtractions between overlapping plate counts. Given the confirmed selectivity and specificity of each agar plate, this provided a quantitative description of the dynamics of each subpopulation throughout the conjugation experiments.

Plating of transconjugants T onto ATBRh, ATBT, and ATBTS agar plates revealed that small proportions of transconjugants was able to grow on ATBRh (ratio_R_T = 6.02·10^-2^) and ATBTS (ratio_TS_T = 5.37·10^-7^). Results per plated colony are presented in Supp. material S2 and estimated ratios are presented in Supp. material S3.

Plating of recusants and escapers confirmed that growth was strictly restricted to the expected selective media for each subpopulation, with no growth observed on any other agar tested (Supp. material S4 and Supp. material S5).

For the analysis of data, the five different subpopulation concentrations were calculated from enumerations on antibiotic-containing agar plates by solving the following system of equations (1):

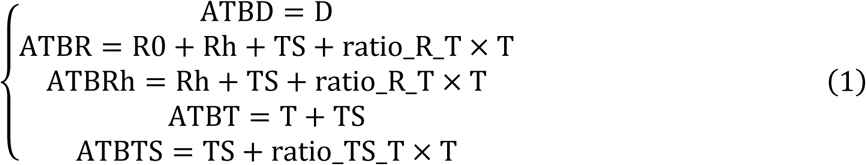

where ATBD, ATBR, ATBRh, ATBT and ATBTS denote the total bacterial concentrations enumerated on their respective antibiotic-containing agar plates; and D, R0, Rh, T and TS denote the concentrations of donors, recipients, recusants, transconjugants and escapers respectively.

By comparing subpopulation proportions after 8 h of conjugation with multiplex PCR (Supp. material S1.) results obtained at the same time point, we could show that the transconjugants recovered on ATBRh had all lost the TAP, indicating that this residual growth reflects TAP loss rather than helper-only transfer.

#### 1.2 Growth kinetics

To characterize the intrinsic growth properties of each subpopulation, growth kinetics were monitored for donors, recipients, transconjugants, and escapers isolated from conjugation experiments. As shown in Figure 2, all populations displayed typical bacterial growth dynamics, with a distinct exponential phase followed by a transition into stationary phase, reaching approximately 10^9^ CFU·mL^-1^ over the 24h period. Growth rates were broadly comparable across all subpopulations, although donors appeared to grow slightly faster. Bacterial concentrations on all antibiotic-containing agar plates reached approximately 10^8^ to 10^9^ CFU·mL^-1^ by 24 h.

**Figure 2.**
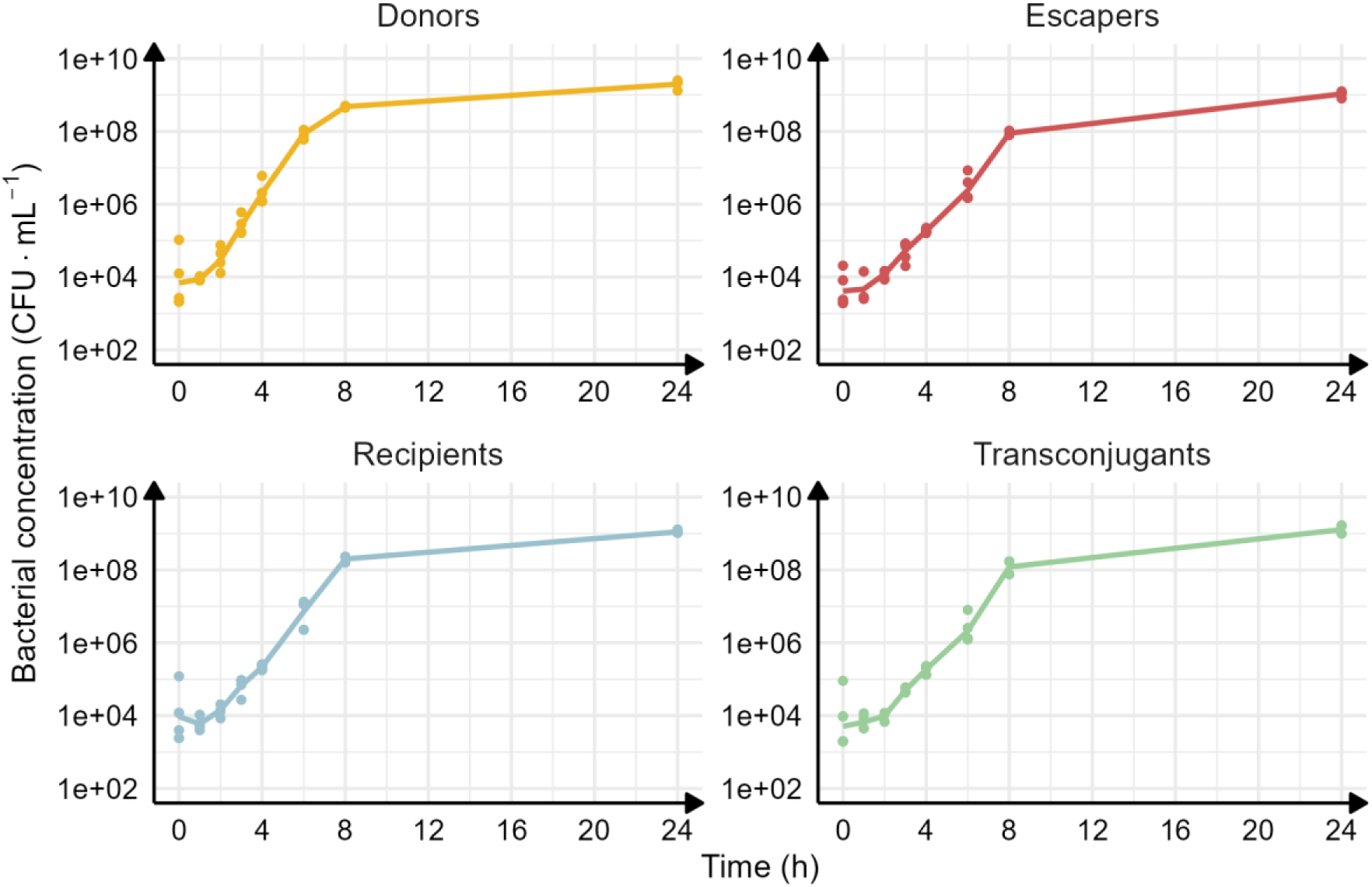
Growth kinetics of bacterial populations involved in conjugation. Average growth curves for donors, recipients, transconjugants, and escapers are shown, with individual replicate measurements (initial inoculum: 10^4^ CFU·mL^-1^; n = 3 for transconjugants, n = 4 for the other subpopulations)

#### 1.3 Conjugation kinetics

A total of 44 conjugation time-course experiments were performed across different donor-to-recipient ratios (1:100, 1:3, 3:1 and 100:1), initial inocula (10^2^, 10^4^, 10^6^ and 10^8^ CFU·mL^-1^), and growth phases (stationary or exponential). As an example, Figure 3 shows enumeration results for the typical conditions: initial bacterial inoculum in stationary phase with a starting inoculum of 10^6^ CFU.mL^-1^ and a D:R ratio of 1:100 (n=4). All replicates showed consistent growth and conjugation dynamics, demonstrating the reproducibility of the measurements and reliability of the experimental setup.

**Figure 3.**
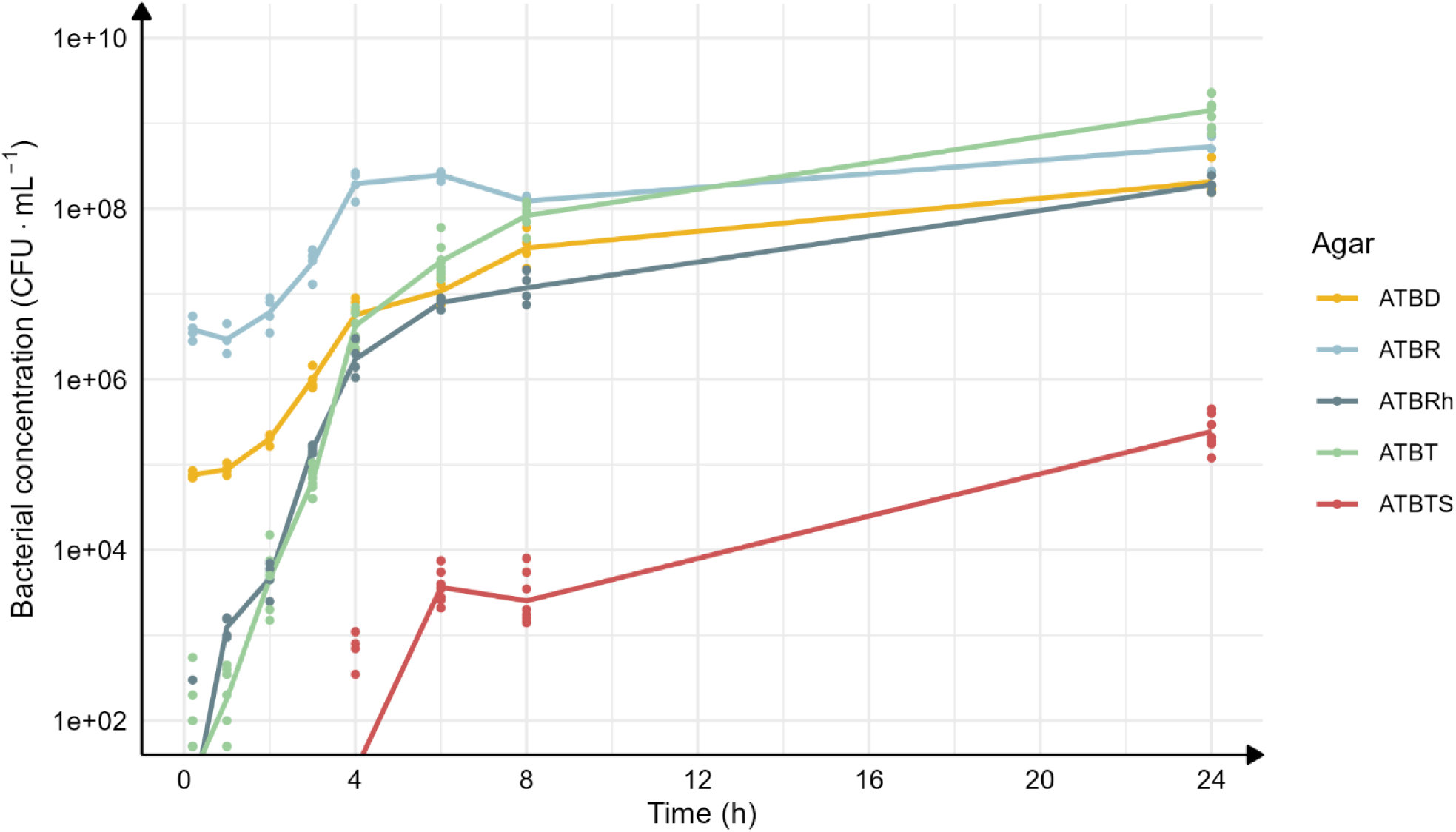
Example of conjugation dynamics. Conjugation dynamics are shown for a representative condition with a starting inoculum of 10^6^ CFU·mL^-1^ in stationary phase and a D:R ratio of 1:100. Each curve represents mean viable cell counts obtained from each antibiotic-containing agar plates (n = 4 for ATBD, ATBR and ATBRh, n = 8 for ATBT and ATBTS).

Colonies on ATBD and ATBR were detectable from the earliest time points, increased exponentially, and reached stationary-phase concentrations plateau of approximately 10^8^–10^9^ CFU·mL^-1^ around 10– 12 h. Colonies on ATBT were absent at t = 0 but subsequently increased exponentially and reached a similar plateau by10–12 h, consistent with the progressive emergence of helper-carrying bacteria following conjugation. Colonies on ATBRh were also detectable early but exhibited slower initial dynamics, with a marked increase occurring after approximately 4 h. In contrast, colonies on ATBTS emerged later, becoming detectable only after approximately 4–5 h, and remained substantially less abundant throughout the experiment. At 24 h, their concentration reached approximately 10^5^ CFU·mL^-1^ at 24 h) several orders of magnitude below those measured on the other selective media (ATBD, ATBR, ATBRh and ATBT).

Growth and conjugation data were used together to develop and calibrate the mathematical model described below. Data are available at https://doi.org/10.57745/SZGWPL.

### 2. Modeling

#### 2.1 Tested hypotheses

The ability of transconjugants to act as secondary donors was investigated. Since transconjugants receive the helper plasmid upon TAP transfer, they may themselves initiate conjugation with remaining recipients. This hypothesis was supported by the data and retained, introducing the transconjugant-mediated conjugation rates γ_T_ and γ_TTS_.

The origin of escapers (TS) was also tested. Two possible routes were considered: emergence *via* conjugation from donors or transconjugants (rates γ_DTS_ and γ_TTS_), or direct transition from the transconjugant population independently of conjugation. Only the conjugation-driven routes were supported by the data and retained in the final model, indicating that escapers arise exclusively through conjugation events, from both donor and transconjugant populations.

The origin of recusants (Rh) was investigated through alternative formation pathways. Their generation from R0 *via* donor-mediated conjugation, occurring at rate γ_Dh_, was supported by the data and therefore retained in the model. In contrast, conjugation mediated by transconjugants was not supported and was excluded. Additionally, the parameter k_TR_ was included to account for TAP loss from transconjugants (T) leading to the formation of recusant clones.

The hypotheses tested for escaper formation considered their potential origin *via* conjugation with donors, transconjugants, and/or escapers. Only the first two pathways, involving donors and transconjugants, were supported by the data and retained in the model, whereas the contribution from TS was excluded.

Other tested but rejected hypotheses are illustrated in Supp. material S6.

#### 2.2 Final model

The model is based on a system of ordinary differential equations (ODEs) describing the temporal dynamics of five populations: donors (D), recipients (R0), recusants (Rh), transconjugants (T), and escapers (TS) (Supp. material S7.). The full structure of the model is illustrated in Figure 4.

**Figure 4.**
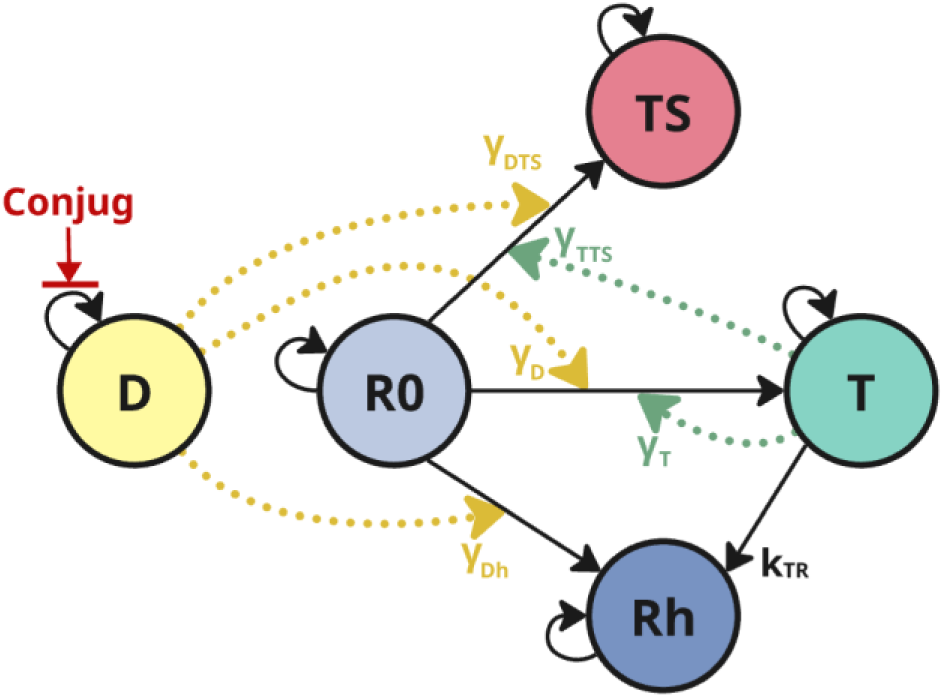
Schematic representation of the mathematical model describing TAP conjugation dynamics. Five populations are modeled: donors (D), recipients (R0), transconjugants (T), escapers (TS) and recusants (Rh). Solid black arrows represent conjugation-driven transitions from R0 to T, TS or Rh. Dotted yellow arrows indicate donor-mediated conjugation rates (γ_D_, γ_DTS_, γ_Dh_), while dotted green arrows indicate transconjugant-mediated conjugation rates (γ_T_, γ_TTS_). The parameter k_TR_ denotes the transition rate from T to Rh (TAP loss). Self-looping arrows indicate population growth. The red arrow indicates reduction of donor growth rate under conjugation conditions.

Bacterial growth was modeled using logistic growth for all populations. Conjugation processes, corresponding to TAP transfer, were modeled as mass-action interactions driven by both donor and transconjugant populations.

#### 2.3 Model fit

Figure 4 provides a schematic overview of the model structure; the underlying equations describe the dynamics of the five subpopulations (donors, recipients, transconjugants, recusants, and escapers). Model performance was evaluated using visual predictive checks (VPCs), demonstrating good agreement between observed and model-predicted data across all experimental conditions.

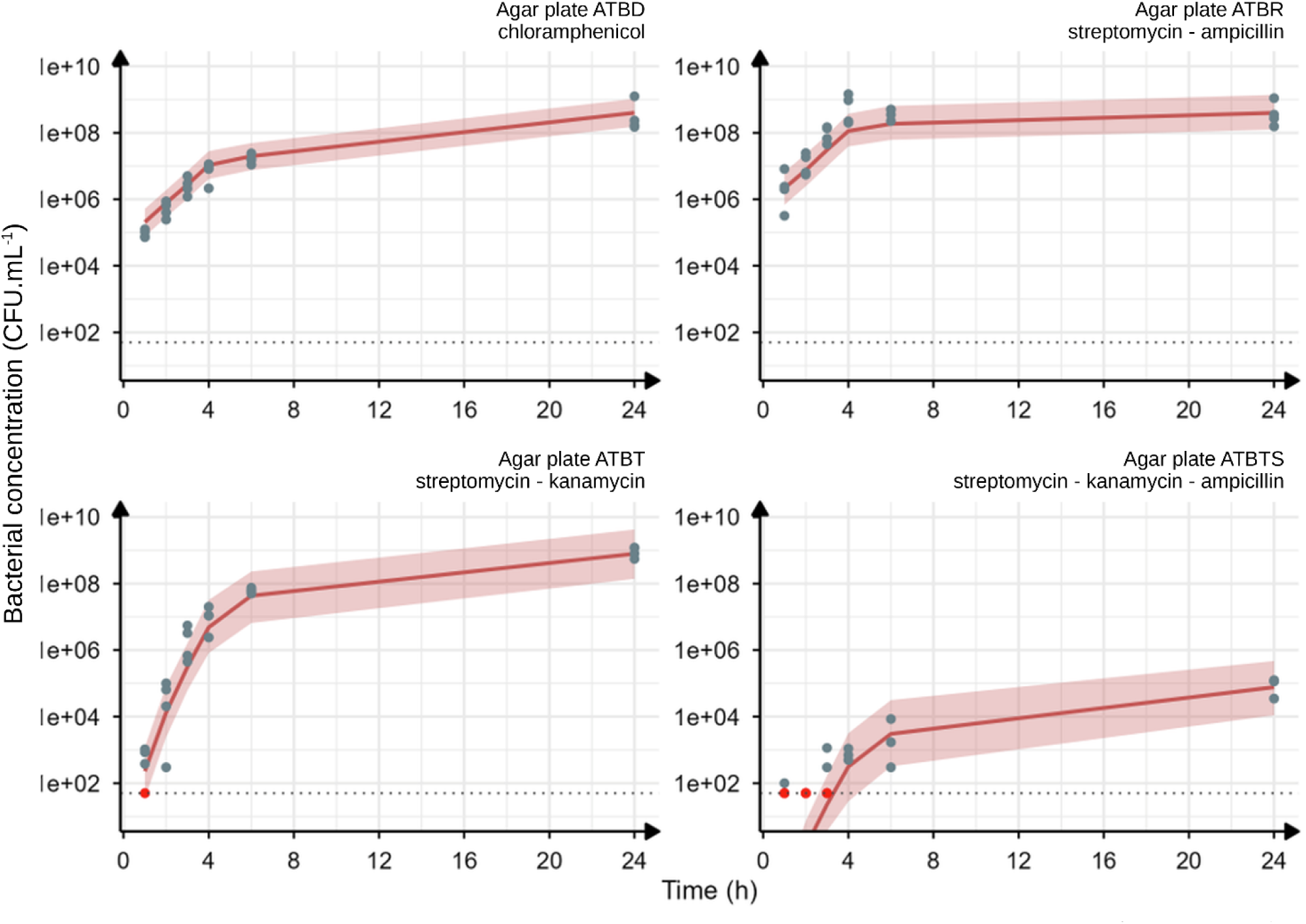

Figure 5 shows VPC results for a representative condition (initial inoculum of 10^6^ CFU·mL^-1^ in stationary phase, D:R ratio of 1:100); VPCs for all remaining conditions are provided in the Supplementary Materials (Supp. material S8.). Individual predicted versus observed profiles were additionally inspected visually for each experiment and were found consistent with the VPCs, confirming the model’s ability to adequately describe individual bacterial dynamics across the full range of conditions tested (Individual fits and observations vs predictions provided in Supp. material S9. and S10.).

**Figure 5.**
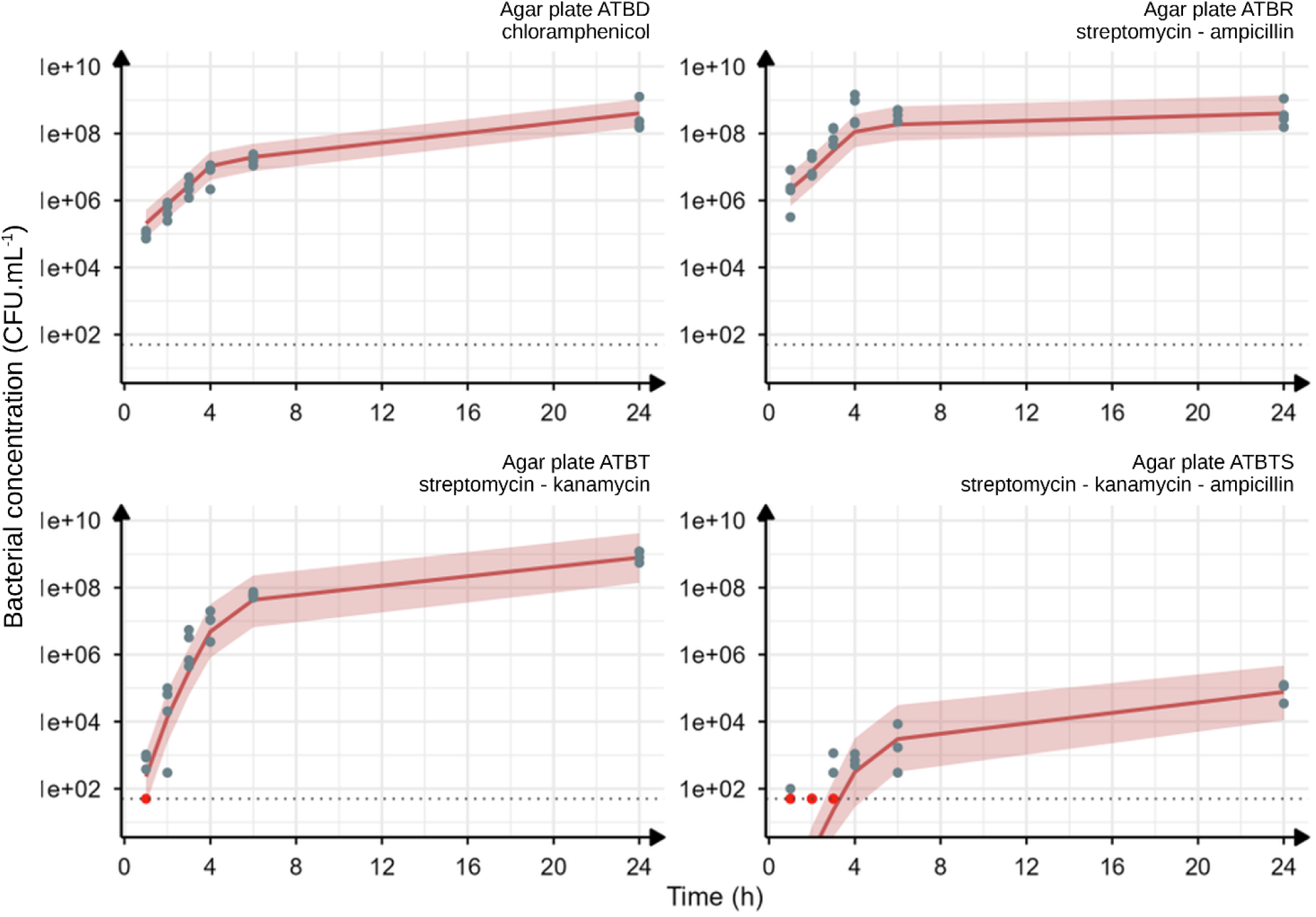
Visual predictive check (VPC) of the final model for a donor inoculum of 10^6^ CFU·mL^-1^ and a D:R ratio of 1:100. Blue dots represent observed data, red dots indicate values below the limit of quantification (50 CFU.mL^-1^ - dotted line). The red line shows the median prediction, and the shaded area represents the 90% prediction interval (5^th^–95^th^ percentiles). Panels correspond to each antibiotic-containing agar: ATBD (chloramphenicol), ATBR (streptomycin and ampicillin), ATBT (streptomycin and kanamycin), and ATBTS (streptomycin, kanamycin, and ampicillin). Results are based on 500 simulations.

#### 2.4 Parameters

##### 2.4.1. Parameter estimation

Estimated model parameters are reported in Supp. material S1. Overall, parameter estimates showed good precision, with relative standard errors below 10% for most parameters, indicating robust parameter identifiability.

Estimated doubling times were comparable between donor and non-donor populations, with a slightly shorter value observed for donors (0.43 h, 95% CI [0.41, 0.45]) than for non-donors (0.46 h, 95% CI [0.44, 0.48]). Despite this small difference, two distinct doubling times were retained in the final model rather than a single shared value, as this improved the OFV and resulted in better VPCs.

Regarding the origin of recusants, comparison of the two mechanisms suggests that recusants primarily originate from partial conjugation events involving donors (γ_Dh_), where only the helper plasmid is transferred (90 to 98% depending on the scenario), while TAP loss in transconjugants (k_TR_) constitutes a secondary pathway (2 to 10%).

Only one parameter exceeded the 30% relative standard error (RSE) threshold classically used to validate parameter precision in population models: the lag time for the donor population (λD, RSE = 38.1%), which likely reflects the limited information available in the experimental data to precisely identify this process. Residual errors ranged from 0.46 log_10_(CFU·mL^-1^) (donors) to 0.92 log_10_ (escapers), all remaining below the 1 log_10_(CFU·mL^-1^) acceptability threshold. The higher residual variability observed for the escaper population (aATBTS = 0.92 log_10_(CFU·mL^-1^)) is consistent with the rarity of this subpopulation, which leads to greater stochasticity in counts on antibiotic-containing agar plates.

##### 2.4.2. Conjugation dynamics

Conjugation rate estimates revealed distinct contributions of donor and transconjugant populations to TAP transfer. The conjugation rate from donors γ_D_ was more than twenty times higher than that from transconjugants γ_T_ (Supp. material S11).

Additional conjugation pathways leading to escapers (TS) and recusants (Rh) were associated with lower transfer rates, reflecting the lower frequency of these transitions (γ_DTS_, γ_TTS_ and γ_Dh_; values on Supp. material S1).

##### 2.4.3. Growth modulation

The estimated transition rate from transconjugants to recusants (k_TR_) indicates TAP loss. Growth modulation parameters further revealed a reduction in growth at high population densities, with a threshold effect captured by B_50_ and a strong reduction factor (β = 0.85), consistent with density-dependent limitations.

A growth penalty was also observed in donor populations during conjugation experiments compared to growth kinetics alone (18% decrease), suggesting either a fitness cost associated with plasmid carriage and/or conjugation activity, or competitive interactions between different bacterial strains.

#### 2.5 Simulations

Model-based simulations were performed to explore the dynamics of bacterial subpopulations under varying initial donor-to-recipient ratios and inoculum sizes. An example is shown in Figure 6, for an initial donor concentration of 10⁵ CFU·mL^-1^ and a D:R ratio of 1:100. Under these conditions, the recipient population was almost entirely converted by 24 h, with virtually no recipients remaining.

**Figure 6.**
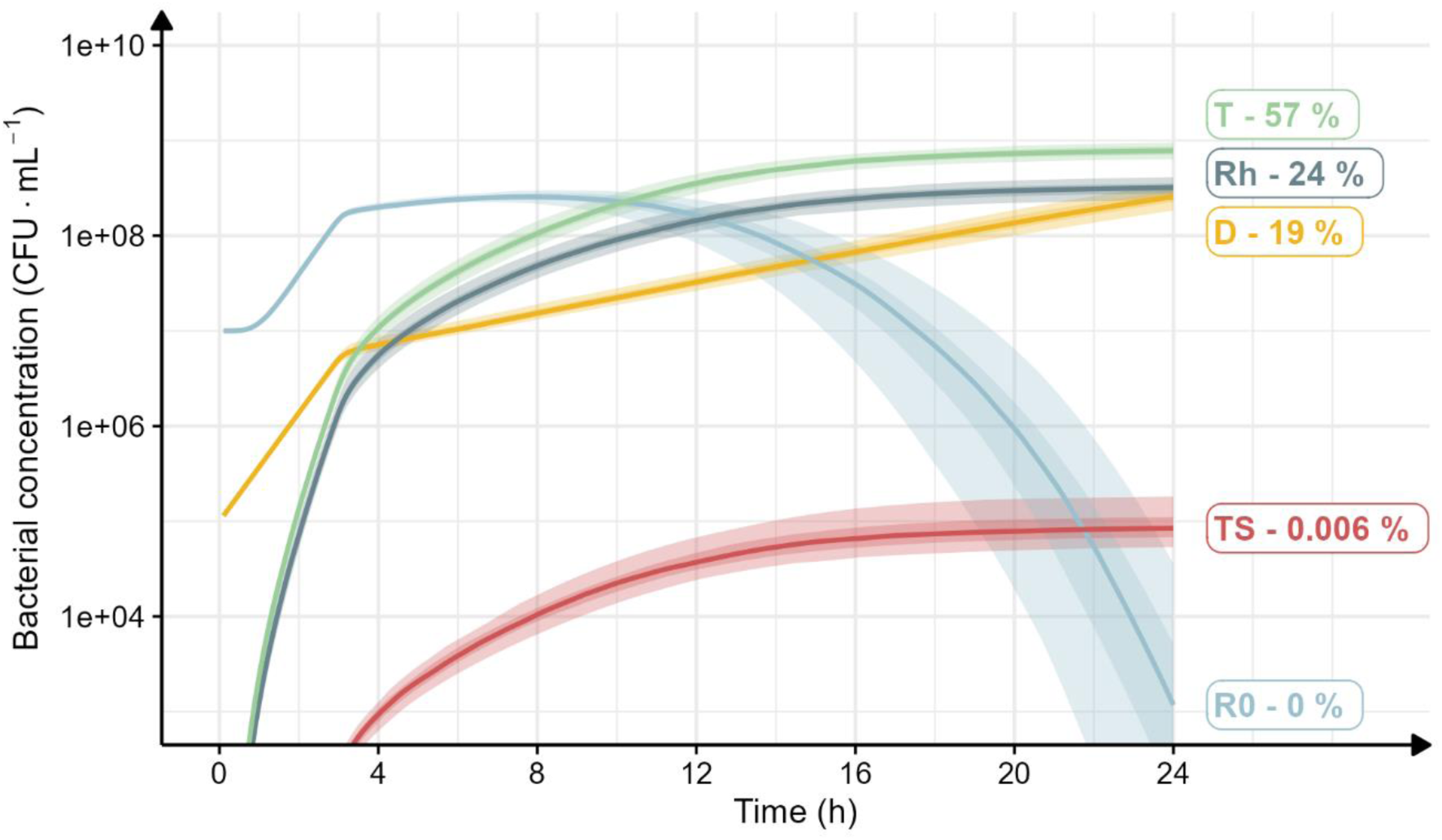
Typical dynamics of bacterial subpopulations during conjugation. Model-predicted dynamics of bacterial subpopulations during conjugation over 24 h. Simulations were performed with initial concentrations of 10^5^ CFU·mL^-1^ donors [D] and a D:R ratio of 1:100. Relative population proportions at 24 h are also shown. Solid lines represent median predictions; shaded areas indicate uncertainty associated with parameter estimation (25^th^-75^th^ and 5^th^-95^th^ percentiles).

Additional simulations performed across the full range of inocula and D:R ratios confirmed that the recipient population consistently decreased substantially, with reductions reaching up to 100% depending on the initial conditions. These results demonstrate that donors have the capacity to contact all target bacteria, as no recipients remained uncontacted after 24 h regardless of the initial experimental conditions. However, not all contacted bacteria received the TAP, as a substantial proportion acquired only the helper plasmid and formed the recusant subpopulation.

### 3. Efficiencies

For a donor inoculum of 10^5^ CFU·mL^-1^ and a D:R ratio of 1:100, more than 70% of the target recipient population was resensitized by TAP transfer and subsequently classified as transconjugants (Figure 7).

**Figure 7.**
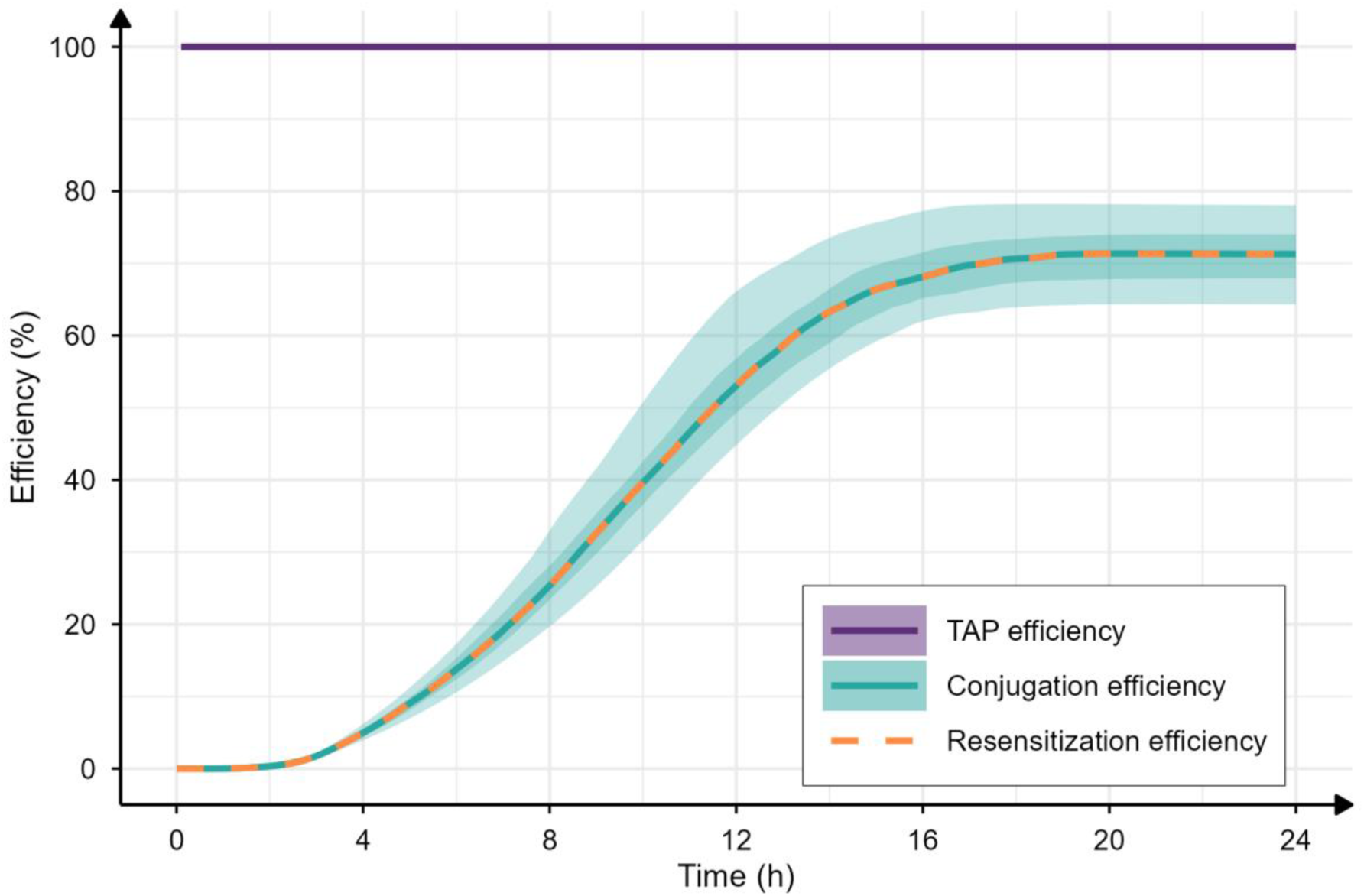
Simulated efficiencies with 10^5^ CFU·mL^-1^ donors and 10^7^ CFU·mL^-1^ recipients [R0]. Model-predicted dynamics of TAP efficiency, conjugation efficiency, and the resulting resensitization efficiency during conjugation over 24 h. Simulations were performed with initial concentrations of 10^5^ CFU·mL^-1^ donors [D] and 10^7^ CFU·mL^-1^ recipients [R0]. Solid lines represent median predictions; shaded areas indicate uncertainty associated with parameter estimation (25^th^-75^th^ and 5^th^-95^th^ percentiles). For resensitization efficiency, only the median prediction is shown (dashed line) for clarity.

Across all simulated conditions, TAP efficiency was nearly fully effective (less than 0.01% of escapers for all simulations), indicating that the limiting factor in resensitization is conjugation efficiency rather than TAP efficiency.

The impact of initial inoculum size on resensitization efficiency was further explored (Figure 8). While the maximal resensitization plateau remained above 70% across conditions, higher inoculum sizes led to a faster achievement of this plateau.

**Figure 8.**
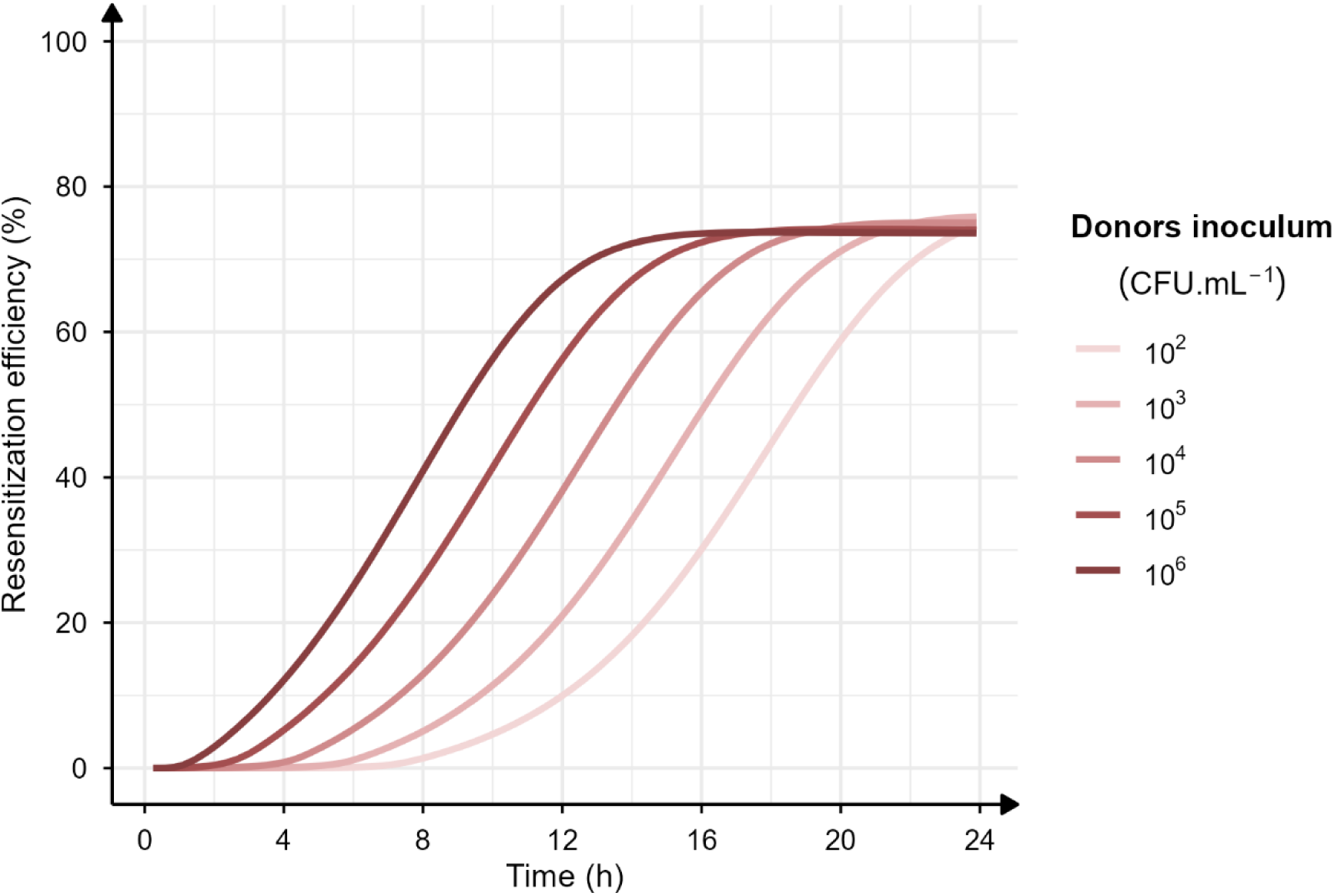
Resensitization efficiencies for a D:R ratio of 1:100 depending of the donor inoculum. Simulated resensitization efficiency for donor inocula ranging from10^2^ to 10^6^ CFU·mL^-1^.

The influence of the donor-to-recipient ratio was also investigated for a fixed donor inoculum of 10^5^ CFU·mL^-1^ (Figure 9). A moderate increase in resensitization efficiency was observed at lower D:R ratios, yet resensitization consistently plateaued before completion, regardless of the initial experimental conditions.

**Figure 9.**
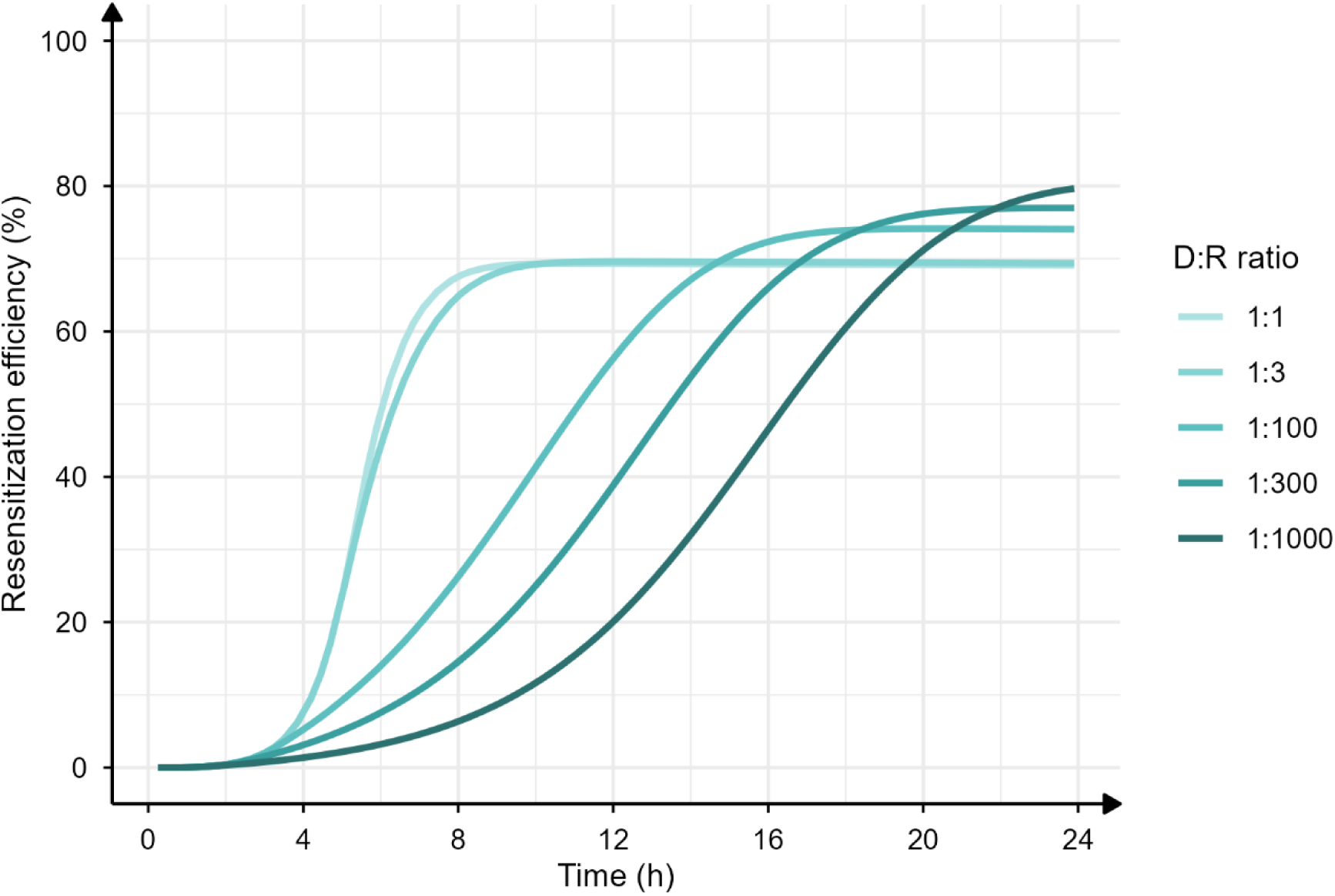
Resensitization efficiencies depending of the Donors:Recipients ratio for a donor inoculum of 10^5^ CFU·mL^-1^. Simulated resensitization efficiency for D:R ratios from 1:1 to 1:1000.

## Discussion

In this study, we combined experimental conjugation data with mathematical modeling to evaluate the transfer dynamics of TAP_F_-dCas9-OXA48 and its ability to resensitize resistant bacteria, *i.e.*, the capacity of the CRISPR/dCas9 system to inhibit expression of the *bla*_OXA-48_ gene carried on pOXA-48, which confers ampicillin resistance in recipients. Our results provide a quantitative framework for understanding the potential and limitations of this strategy, identifying key parameters governing its efficiency, and offering guidance for optimizing TAP deployment against antimicrobial resistance.

### 1. Longitudinal modeling identifies delivery as the limiting step

In most conjugation studies, transfer efficiency is typically estimated by single end-point measurements (*e.g.*, ranging from 2 hours to overnight (11,12)), capturing only a static snapshot of a fundamentally dynamic process. In contrast, we assessed it over a 24 h conjugation period using a longitudinal approach, made possible by explicitly modeling the true bacterial subpopulations rather than raw colony counts on antibiotic-containing agar plates; a distinction required because plating itself introduces a systematic bias, as transconjugants that had lost the TAP plasmid, or that had escaped CRISPR/dCas9-mediated repression, could revert to a resistant phenotype in the timeframe between plating and colony enumeration. Correcting this bias revealed four concurrent dynamics: the sustained presence of donors (D) actively participating in conjugation at late time points; the marked decline of the recipient (R0) population, concomitant with an increase in transconjugants (T); the low but steady presence of escapers (TS); and the progressive emergence of recusants (Rh).

Across all simulated scenarios, TAP efficiency (*i.e.*, the ability of the CRISPR/dCas9 system to resensitize a recipient once the TAP has been acquired) was estimated at effectively 100% throughout the 24 h period. This indicates that resensitization is not limited by the biological activity of the CRISPR/dCas9 system itself, but rather by its delivery: overall resensitization efficiency mirrors conjugation efficiency almost exactly, owing to the very low and stable emergence of escapers (TS/T ratio never exceeding 10^-4^ across all scenarios, consistent with previously reported frequencies (6,13)). In other words, the bottleneck of resensitization lies not in target inhibition, but in getting the construct into the target cell in the first place.

### 2. Determinants of resensitization efficiency

This decomposition of efficiency into elementary components (conjugation efficiency, CRISPR/dCas9-mediated resensitization efficiency, and their combined overall efficiency) offers a more general framework for evaluating and comparing conjugation-based antibacterial strategies. By disentangling TAP delivery from CRISPR/dCas9 activity, this framework enables identification of the step that most strongly limits overall efficiency and should therefore be prioritized for optimization. This provides greater mechanistic insight than a single aggregate efficiency metric, which may obscure distinct or opposing processes contributing to the final outcome.

Simulations conducted across different inoculum levels and D:R ratios further show that neither increasing the inoculum nor shifting the D ratio in favor of donors was sufficient, by itself, to overcome the observed resensitization plateau. These findings suggest that the plateau cannot be explained solely by insufficient donor availability or bacterial density and instead point toward recusant formation as an intrinsic constraint on TAP dissemination. Understanding the mechanisms governing the emergence of this refractory subpopulation is therefore critical for identifying strategies to overcome this bottleneck.

### 3. Helper-only transfer creates a refractory population

The main factor limiting overall resensitization efficiency identified in this study is the formation of recusants (clones that have received the helper plasmid alone, without the TAP construct). Because the helper plasmid encodes exclusion proteins, its transfer independently of the TAP renders these cells refractory to any subsequent conjugation attempt, effectively removing them from the pool of resensitizable recipients. This dissociated, incomplete transfer of the two-plasmid system constitutes a direct consequence of the *trans* configuration.

This phenomenon was reported by Reuter *et al*., who observed that resensitization of a pOXA-48-carrying recipient population plateaued after 24 h of conjugation with a TAP donor, with a stable ∼90% of recipients becoming ampicillin-susceptible while the remaining ∼10% persisted as resistant (5). Using plasmid profiling at later time points, the authors showed that this residual resistant fraction consistently carried the F-Tn10 helper plasmid without the TAP, confirming that acquisition of the helper alone, and the resulting establishment of F-encoded exclusion, rendered these cells permanently refractory to subsequent TAP acquisition. They further proposed that deleting the helper’s origin of transfer could prevent this outcome, at the cost of a slower, but potentially more complete, dissemination of the TAP across the recipient population.

This vulnerability is structural, and comparing our system to that studied by Hamilton *et al*. helps explain why. They compared a self-transmissible, conjugative plasmid (*cis* configuration) to a mobilizable plasmid relying on a separate, non-self-transmissible helper lacking an *oriT* (*trans* configuration) (14). In their *cis* system, transconjugants themselves became new donors, driving an exponential increase in conjugation frequency over time. Their *trans* system showed no such amplification, precisely because the helper, lacking an *oriT*, was never co-transferred with the mobilizable plasmid, leaving transconjugants unable to conjugate further.

Our system differs in one key respect: unlike Hamilton’s *trans* configuration, our helper plasmid does carry an *oriT* and is therefore co-transferred alongside the TAP. As a result, our transconjugants can act as secondary donors, like in a *cis* configuration. However, this same co-transfer mechanism is also what allows the helper to occasionally disseminate on its own, independently of the TAP, generating recusants. This is precisely the trade-off underlying the *oriT*⁻ helper strategy discussed above: removing the helper’s *oriT*, as in Hamilton’s *trans* system, would eliminate recusant formation, but at the likely cost of losing this secondary-donor effect, and therefore of a lower overall conjugation frequency.

### 4. Implications of the findings: design rules for next-generation TAP delivery

Since resensitization efficiency is ultimately governed by conjugation efficiency, and recusant formation *via* independent helper transfer is its principal limiting factor, improving TAP-based systems should focus primarily on eliminating this dissociated transfer, rather than on further optimizing CRISPR/dCas9 activity itself, which is already close to maximal.

One direct solution, use an *oriT*-deficient helper plasmid (13): a helper capable of mobilizing the TAP in *trans* but incapable of self-transfer in the absence of its own origin of transfer. Under this design, the helper can no longer disseminate independently of the TAP, which would abolish both the recusant subpopulation and the associated exclusion phenomenon altogether.

To illustrate the magnitude of this potential gain, we ran a simulation from the final model in which the helper plasmid’s transfer parameters were set to zero, *i.e.*, the helper is no longer transmitted on its own, and transconjugants can no longer act as secondary donors of the helper alone. Under these conditions, resensitization efficiency increased up to 100% in most scenarios (see Supp. material S12.); in the remaining scenarios, starting from a small donor and/or recipients’ inoculum, maximum efficiency was not yet reached within the 24 h simulation window. This result should, however, be interpreted with caution: it does not constitute a validated simulation of an *oriT*-deficient helper system, since the experimental data used to build the model were generated with a fully transferable helper plasmid, and the model was not calibrated to describe the behavior of an *oriT*-deficient helper. It should therefore be read as an illustrative upper-bound projection of what eliminating helper-only transfer could achieve, rather than a quantitative prediction.

Beyond the *oriT*-deficient helper strategy, several complementary design routes could be considered. Addressing the helper-driven exclusion problem itself, a single-plasmid, *cis* configuration integrating both the CRISPR/dCas9 system and the conjugation machinery would remove the dependency on a separately transferred helper altogether (14), at the cost of increased construct complexity. Separately, to further limit escaper emergence, targeting *bla*OXA-48 with multiple guide RNAs would not be expected to substantially reduce escaper frequency, since escapers appear to arise predominantly from inactivation of Cas9 itself, *e.g. via* transposon insertion, rather than from mutation or loss of the target locus. A more promising strategy would instead rely on delivering two independent TAP constructs: since escaper emergence would then require independent Cas9 inactivation events on both constructs, the resulting escaper frequency would be expected to scale as the square of the single-TAP inactivation frequency (f²), rather than f, offering a substantially greater reduction in escaper emergence (13).

Alternatively, conjugation-free delivery systems, such as bacteriophages (15,16) or nanoparticle-based vectors (17), could be explored.

In conclusion, we developed a mathematical model integrating experimental data to describe TAP-mediated bacterial resensitization *via* conjugation. While TAPs are highly effective at resensitizing recipients once acquired (∼100%), overall resensitization efficiency (up to 80%) is limited by TAP transfer dynamics (specifically, the independent transfer of the helper plasmid, which generates recusants refractory to further conjugation). Removing this bottleneck, whether through an *oriT*-deficient helper design or alternative delivery architectures, represents the clearest lever for improving TAP performance, and underscores the value of longitudinal, mechanistic modeling for guiding the rational design of next-generation antimicrobial resistance strategies.

## Materials and methods

### 1. Experiments

#### 1.1 Bacterial strains, plasmids, antibiotics and medium

##### 1.1.1. Bacterial strains and plasmids

Experiments were performed using two *Escherichia coli* genotypes. Genotype K12 F- λ- *rph-1* DE(*lacIZYA)::FRT* (lac-) was used as the donor (D); it carries a chromosomal chloramphenicol resistance marker, TAP (carrying a kanamycin resistance gene) and a helper plasmid F (carrying a tetracycline resistance gene). Genotype K-12 MG1655 *rpsL* was used as the recipient (R0); it carries a Spontaneous mutation to streptomycin resistance marker and a pOXA-48 plasmid. The different resistance genes allow selective tracking of each sub-populations in mixed cultures.

The pOXA-48 plasmid is a natural plasmid carrying a *bla*_OXA-48_ gene, which confers resistance to β-lactam antibiotics, including carbapenems and ampicillin. Ampicillin was used in this study as a selection marker to track the presence of pOXA-48 plasmid.

The TAP (TAP_F_-dCas9-OXA48) is based on a CRISPR/dCas9 plasmid encoding the complete CRISPR/dCas9 system with a single guide RNA targeting the promoter of the *bla*_OXA-48_ gene (5), thereby mediating transcriptional repression of this resistance gene. A kanamycin resistance gene is included on the plasmid as a selectable marker, enabling selection of TAP-carrying bacteria on antibiotic-containing agar plates.

##### 1.1.2. Media

Bacteria were grown in Lysogeny Broth medium (Sigma-Aldrich, Saint-Quentin-Fallavier, France) at 37°C. Subpopulation selections were performed on antibiotic-containing LB agar plates as described in Table 2. The antibiotics used were: ampicillin (Carl Roth, Lauterbourg, France; cat. no. K029); kanamycin (Sigma-Aldrich; cat. no. 60615); tetracycline (Sigma-Aldrich; cat. no. PHR1041); chloramphenicol (Sigma-Aldrich; cat. no. C0378); streptomycin (Sigma-Aldrich; cat. no. S6501). Antibiotic concentrations in antibiotic-containing agar were adopted from Reuter *et al.* (5).

#### 1.2 Identification and enumeration of bacterial subpopulations

Conjugation was performed between **donor** bacteria (**D**) and the initial **recipient** bacteria (**R0**).

Following conjugation, recipient-derived subpopulations were identified and enumerated based on their plasmid content and expression, as inferred from their antibiotic resistance phenotype:

- **Rh**: **Recusants**, defined as recipient-derived bacteria harboring the helper plasmid only (refractory for further conjugation);
- **T**: **Transconjugants**, defined as recipient-derived bacteria that acquired TAP (and helper plasmid) and were resensitized to ampicillin through active CRISPR/dCas9-mediated repression;
- **TS**: **Escapers**, defined as TAP and helper-containing recipient-derived bacteria that remained ampicillin-resistant due to escape from CRISPR/dCas9-mediated repression.

Table 1 summarizes the characteristics of each subpopulation identified in the conjugation experiments.

**Table 1.**
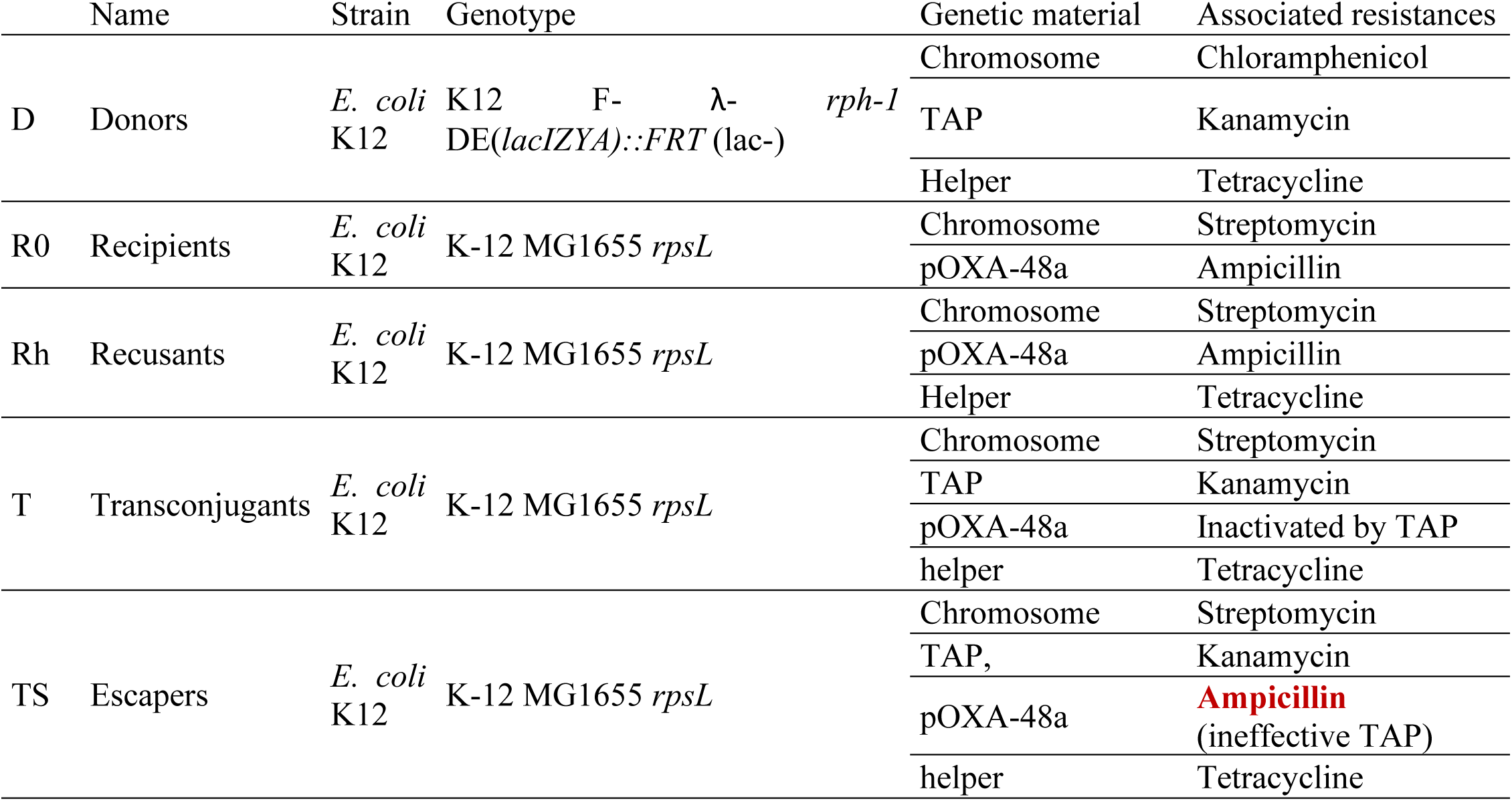
Bacterial strains and their associated plasmid content and resistance profiles. Bacterial subpopulations used in conjugation experiments, with their plasmid content and antibiotic resistance phenotypes and the antibiotic-containing agar supporting their growth. Ampicillin resistance, targeted by TAP, is highlighted in red.

Bacterial subpopulations were quantified based on their phenotypic characteristics (antibiotic resistances) and their growth on antibiotic-containing agar plates. Each bacterial population was either evaluated directly by counting the number of CFU on the corresponding antibiotic-containing agar plate or by calculating the difference in the number of CFU between two complementary antibiotic-containing agar plates (see Table 2). Prior validation confirmed that these phenotypic markers reliably reflected the presence or absence of the corresponding plasmids, enabling the correct assignment of bacteria to their respective subpopulations. To confirm the selectivity of each antibiotic-containing agar, 96 clones were isolated from each subpopulation (donors, recipients, recusants, transconjugants, and escapers) and characterized by multiplex PCR (see Supp. Material S1.) to determine the prevalence of each plasmid (TAP, helper, and pOXA-48).

**Table 2.** Composition and selectivity of the agar media. Antibiotic-containing agar employed in this study, detailing the antibiotics incorporated into each formulation, their final concentrations, and the corresponding bacterial subpopulations isolated on each of them. Subpopulations are described in material and methods section 1.2.

| Name attributed to antibiotic-containing agar | Antibiotics | Concentrations | Selected populations |
| --- | --- | --- | --- |
| ATBD | Chloramphenicol (CHL) | 20 mg·L<sup>-1</sup> | D |
| ATBR | Streptomycin (STR) | 20 mg·L<sup>-1</sup> | R0, Rh and TS |
|  | Ampicillin (AMP) | 100 mg·L<sup>-1</sup> |  |
| ATBRh | Streptomycin (STR) | 20 mg·L<sup>-1</sup> | Rh and TS |
|  | Ampicillin (AMP) | 50 mg·L<sup>-1</sup> |  |
|  | Tetracycline (TET) | 10 mg·L<sup>-1</sup> |  |
| ATBT | Streptomycin (STR) | 20 mg·L<sup>-1</sup> | T and TS |
|  | Kanamycin (KAN) | 50 mg·L<sup>-1</sup> |  |
| ATBTS | Streptomycin (STR) | 20 mg·L<sup>-1</sup> | TS |
|  | Kanamycin (KAN) | 50 mg·L<sup>-1</sup> |  |
|  | Ampicillin (AMP) | 100 mg·L<sup>-1</sup> |  |
| LB agar | - | - | D, R0, Rh, T and TS |

Bacterial concentrations of subpopulations were estimated during experiments (see 1.4. Growth kinetics and 1.5. Conjugation kinetics) by spotting 10 µL twice of serially diluted conjugation mixtures (undiluted to 10^-7^) onto all antibiotic-containing agar plates (ATBD, ATBR, ATBRh, ATBT, and ATBTS; described in Table 2). Plates were incubated at 37°C for 14-18 h, after which colony-forming units (CFU) were enumerated using a SCAN300 colony counter (version 8.6.8.0 v3.4). Bacterial concentrations were calculated as CFU·mL^-1^. antibiotic-containing agar plates

#### 1.3 Antibiotic-containing agar specificity

To verify that subpopulations Rh, T, and TS grew exclusively on their expected selective media (as defined in Table 2), isolated colonies were resuspended in 150 µL of LB broth, prior to plating on the three selective agars (ATBRh, ATBT and ATBTS). Each collected colony was assumed to be phenotypically pure, as confirmed by multiplex PCR (Supp. material S1.).

The procedure was repeated with sixty colonies for transconjugants, previously selected on ATBT. For recusants (selected on ATBRh) and escapers (selected on ATBTS), only twenty-four colonies were selected. The resulting counts were expressed in CFU·mL.⁻¹.

When growth was observed on an agar other than the expected one, the proportion of colonies growing on that unexpected agar was calculated relative to the number of colonies growing on the expected agar. For transconjugants, these proportions, corresponding to ratio_R_T (unexpected growth on ATBRh) and ratio_TS_T (unexpected growth on ATBTS) relative to growth on ATBT, were estimated using the M3 method implemented in Monolix (Monolix 2024R1, Simulations Plus, doi: 10.5281/zenodo.11401936) to handle values below the limit of quantification (LOQ = 50 CFU·mL^-1^).

#### 1.4 Growth kinetics

The growth kinetics of each bacterial subpopulation isolated after a conjugation experiment were monitored under static conditions at 37°C over 24 h (0, 1, 2, 3, 4, 6, 8, and 24 h). Bacterial counts were determined using antibiotic-free agar plates. Experiments were performed with an initial inoculum of approximately 10^4^ CFU·mL^-1^ from both exponential and stationary growth phases.

#### 1.5 Conjugation kinetics

Time-course conjugation assays were performed by mixing donor with recipient bacteria. Bacterial subpopulations were quantified at 0, 1, 2, 3, 4, 6, 8, and 24 h. Experiments were carried out under static conditions at 37°C across a wide range of initial donor inocula (10^2^, 10^4^, 10^6^ and 10^8^ CFU·mL^-1^), donor-to-recipient (D:R) ratios (1:100, 1:3, 3:1 and 100:1), and initial growth phases (exponential or stationary). Enumeration was performed by plating on the different antibiotic-containing agar to quantify each subpopulation as described before.

#### 1.6 Data preprocessing

Growth and conjugation time-course data (bacteria enumerated on antibiotic-containing agar over 24 h) were used to develop a mathematical model comprising a system of nonlinear ordinary differential equations. This framework allows estimation of growth and conjugation-specific parameters, conjugation efficiency, and key factors influencing these processes like initial inoculum or D:R ratio, enabling simulations of various scenarios. Data were formatted for Monolix (Monolix 2024R1, Simulations Plus, doi: 10.5281/zenodo.11401936), log_10_-transformed, and values below the limit of quantification (LOQ = 50 CFU·mL^-1^) were treated using left-censoring.

### 2. Modeling

A mathematical model was developed describing TAP conjugation dynamics together with the growth of the bacterial populations involved.

#### 2.1 Estimation and modelling methods

Model parameters were estimated using the stochastic approximation expectation–maximization (SAEM) algorithm implemented in Monolix (Monolix 2024R1, Simulations Plus, doi: 10.5281/zenodo.11401936). As all experiments were performed under controlled and standardized conditions, no inter-individual variability was included and all parameters were estimated as fixed effects.

Standard errors and log-likelihood were estimated using the linearization method implemented in Monolix.

A constant (additive) error model with normally distributed residuals was used to describe the log_10_-transformed observations error.

Experimental data were formatted and processed using R (18), and the tidyverse package collection (19).

#### 2.2 Tested hypotheses

Several structural models were explored during model development using Monolix (Monolix 2024R1, Simulations Plus, doi: 10.5281/zenodo.11401936). Model selection was guided by the Akaike Information Criterion (AIC), goodness-of-fit diagnostics, visual predictive checks (VPC), and parameter precision (20). A structural hypothesis was retained only if it resulted in a decrease in AIC, acceptable precision of parameter estimates (relative standard error [RSE] ≤ 30%), and biologically plausible parameter values. Exceptions were made for parameters with RSE slightly exceeding this threshold when the parameter was biologically meaningful. Hypotheses were rejected when they did not improve AIC, when RSE exceeded 30%, or when the corresponding parameter estimate was negligible (*i.e*., its confidence interval included 0). Parameter definitions are provided in Supp. material S11.

### 3. Quantification of efficiencies

The overall efficiency of the system, defined as its ability to resensitize recipients (R0), was split into two distinct components. First, conjugation efficiency, corresponding to the system’s capacity to transfer the TAP to recipients. Second, TAP efficiency, defined as the ability of the TAP, once transferred, to restore susceptibility in the recipients. The calculation of these three efficiencies metrics is described below.

#### 3.1 Conjugation efficiency

Conjugation efficiency quantifies the success of TAP transfer to the recipients and is defined as the fraction of recipient-derived bacteria that acquire TAP (Equation 6):

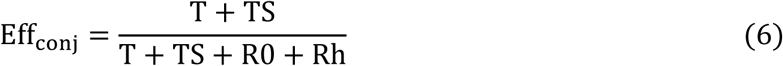

where T, TS, R0 and Rh denote the concentrations of transconjugants, escapers, recipients and of helper-positive recipients, respectively.

#### 3.2 TAP efficiency

TAP efficiency quantifies the intrinsic ability of TAP to resensitize bacteria once transferred and is defined as the fraction of resensitized bacteria among recipient-derived cells harboring TAP (Equation 7):

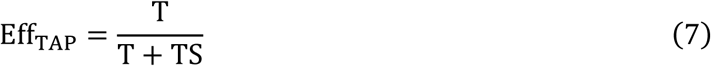

#### 3.3 Resensitization efficiency

Resensitization efficiency quantifies the final biological outcome of the TAP system and is defined as the fraction of recipient-derived bacteria that were successfully resensitized (Equation 8):

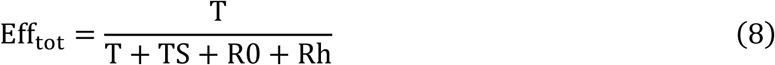

This metric is mathematically equivalent to the product of conjugation efficiency and TAP efficiency (Equation 9):

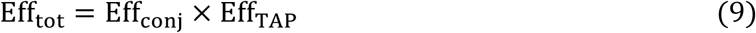

Overall resensitization efficiency constitutes the most biologically relevant metric, as it captures the net resensitization outcome, while intermediate efficiencies enable identification of limiting steps and optimization targets within the TAP system.

## Acknowledgments

We would like to thank Agnès Audurier for her excellent technical assistance.

This work was supported by the ANR through DeCa-P [grant number ANR-22-CE35-0017] and by the Joint Programming Initiative on Antimicrobial Resistance (JPIAMR), under the JPIAMR-ACTION Joint Transnational Call 2021 (grant no. JPIAMR2021-194).

## Data availability

Supplementary materials, including detailed methods and additional figures, are available on Zenodo at https://doi.org/10.5281/zenodo.22178835

